# A non-invasive test for stria vascularis dysfunction

**DOI:** 10.1101/2025.07.18.665193

**Authors:** Neil J. Ingham, Clarisse H. Panganiban, Carolyn M. McClaskey, Kelly C. Harris, Karen P. Steel

**Author notes:** **Author Contributions:** NJI, CHP & KPS designed research. NJI & CHP performed research. NJI, CHP & KCH analyzed data. CCM provided code to process single trial responses. NJI & KPS acquired funding. NJI & KPS wrote the paper. **Competing Interest Statement:** The authors declare no competing interests.

## Abstract

Age-related hearing loss is common in the human population but it is highly heterogeneous in aetiology which has hampered efforts to develop ways of stopping its progression. Three major sites of the initial dysfunction are the sensory hair cells, their innervation, and the stria vascularis which generates the high potassium endolymph maintained at a high endocochlear potential bathing the apical surface of hair cells. Treatments aimed at the initial site-of-lesion may be useful, and diagnostic tools to distinguish the primary site would help stratify clinical trials and facilitate selection of the most suitable treatment for each person. Here we report a new non-invasive test that distinguishes mutant mice with known stria vascularis dysfunction and reduced endocochlear potential from mice with sensory hair cell or neural defects but with normal endocochlear potential. It is based on measuring inter-trial coherence in auditory brainstem responses to individual stimuli. Mice with reduced strial function show good inter-trial coherence compared with the amplitude of the averaged response, while mice with sensory or neural deficits show poor coherence. This method might be useful in humans to predict whether they have reduced strial function, which cannot be measured directly, and identify who would benefit from treatments aimed at boosting strial function.

## Introduction

Hearing impairment is very common in the population and can begin at any age. Childhood deafness affects one in 1000 children born and with age this number increases until over half of adults in their 70s have a significant hearing loss. Hearing impairment isolates people from society, is often associated with depression and cognitive decline, and is a major predictor of dementia (Fellinger et al 2012; Karpa et al 2010; Mick et al 2014; Livingston et al 2017). Hearing aids and cochlear implants can be useful but they do not restore normal function or treat the disease process, so there is a large unmet need for medical approaches to slow down or reverse hearing loss. Recent findings suggest that gene therapy can be successful for children with otoferlin mutations (Lv et al 2024; Qi et al 2024; Wang et al 2024) which is encouraging wider efforts aimed at other genetic types of deafness. However, the vast majority of hearing-impaired people have no genetic diagnosis. Alternative approaches will be needed to widen the scope from targeting specific mutations to treating broad categories of dysfunction.

Histopathology of human temporal bones suggests that age-related, progressive hearing loss predominantly affects three different sites within the cochlea: sensory hair cells; synapses between hair cells and cochlear neurons; and the stria vascularis on the lateral wall of the cochlear duct which generates the high-potassium endolymph maintained at an endocochlear potential (EP) of ∼100mV, essential for normal hair cell function (Schuknecht & Gacek 1993). Any treatment for hearing loss will need to target the initial site of the lesion, so it will be important to be able to diagnose the sites involved. Otoacoustic emissions, generated by outer hair cells, can be used to distinguish a primary inner hair cell/synaptic/neural defect from a primary sensory defect that involves outer hair cells. However, a primary stria vascularis dysfunction is more challenging to distinguish because a reduced EP affects both inner and outer hair cell function and measuring EP directly is an invasive process.

Here we have used a set of mouse mutants with well-characterised primary deficits affecting the three major sites-of-lesion reported in human age-related hearing loss as experimental tools to search for an objective and non-invasive way of differentiating a strial dysfunction from primary sensory or neural defects. Our hypothesis was that in the case of a primary EP reduction, the hair cells may retain some of their normal response properties even when the threshold is raised and the response amplitude is reduced due to a reduced voltage across their transduction channels. In contrast, mice with primary hair cell or neural deficits may show degraded hair cell responses due to their inherent dysfunction. We have discovered that inter-trial coherence in relation to mean Auditory Brainstem Responses (ABR) wave 1 amplitude (ITC/W1amp), recorded using a minimal scalp electrode array at suprathreshold intensities, can distinguish primary strial dysfunction from primary hair cell or neural defects. No other measure used here was able to provide this resolution. The ITC/W1amp measures can be derived from the ABR used clinically. Differential diagnosis will underpin attempts to develop and test treatments for different sites-of-lesion.

## Results

### Mutant mouse panel

A panel of mouse mutants was assembled with hearing loss due to defects at different primary sites-of-lesion within the cochlea, including two with stria vascularis dysfunction and reduced EP (*S1pr2*^*stdf*^ and *Sgms1*^*tm1b*^), one with sensory defects (*Slc26a5*^*tm1*^ affecting outer hair cell function), one with a neural/inner hair cell defect (*Klhl18*^*lowf*^), and three with mixed sensory and neural defects (*Wbp2*^*tm1a*^, *Mir96*^*+14C>A*^ and *Cdh23*^*ahl*^) (Table 1). Each mutant line was analysed at an age when hearing was affected, assessed via ABR thresholds, but there were minimal secondary changes such as hair cell degeneration that would complicate interpretation (Table 1). Littermate controls were analysed alongside the mutant mice at the same ages, and mice from the C57BL/6N line at young ages before the onset of elevated ABR thresholds served as controls for older hearing-impaired C57BL/6N mice, carrying the *Cdh23*^*ahl*^ allele that leads to age-related hearing loss in this inbred strain.

**Table 1.**
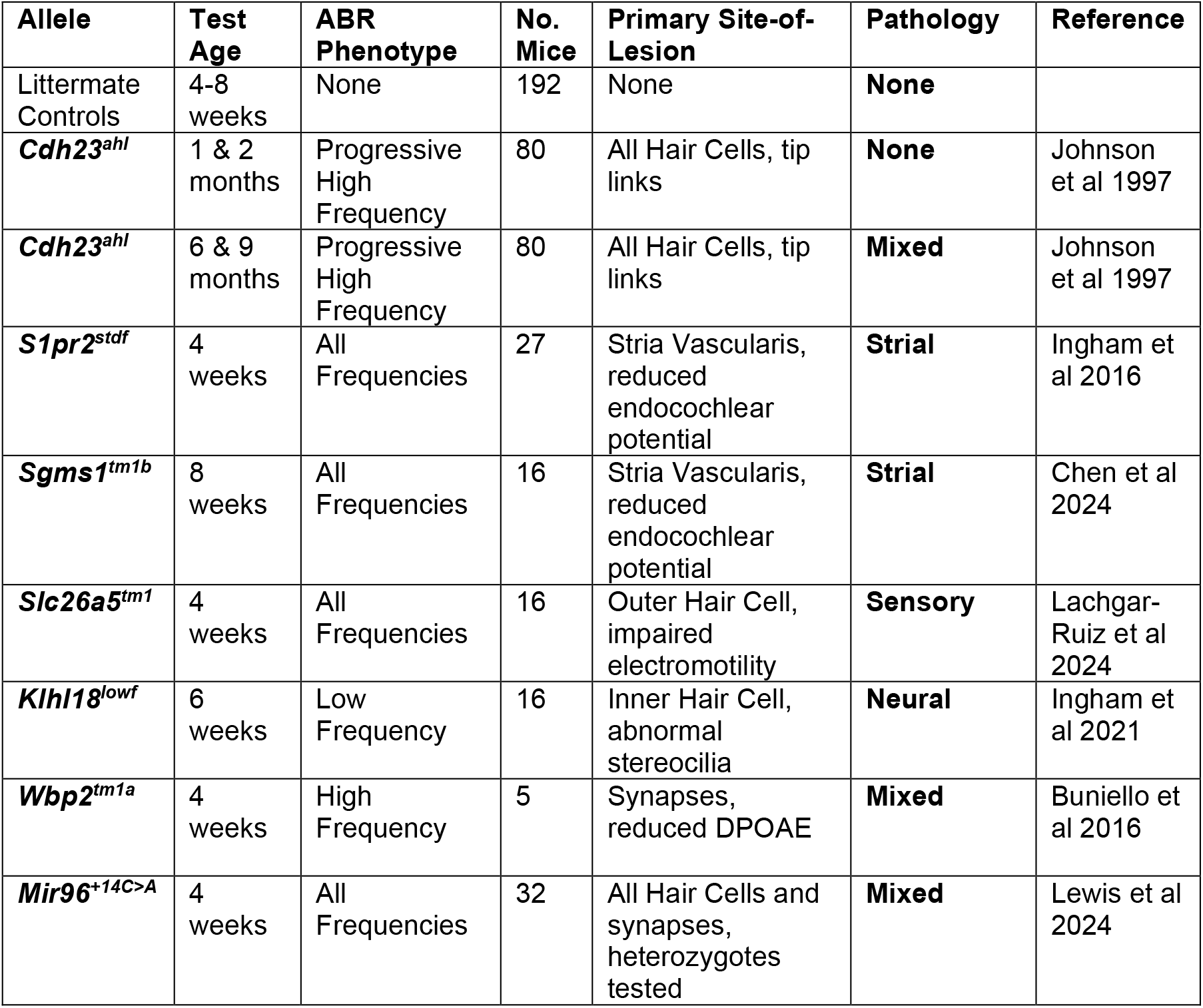
Mutant Mouse lines & associated hearing loss pathology.

### ABR measurements

The thresholds, amplitudes and latencies for the positive peak of ABR wave 1 were measured for three stimulus frequencies (12, 18 and 24 kHz), representing the most sensitive region for hearing in mice. A typical averaged ABR waveform recorded from a train of 512 stimuli at 90 dB SPL is shown in Figure 1 (lower panel). None of these standard ABR measures, or combinations of these measures, allowed any differentiation between the various pathologies in the mutant panel.

**Figure 1.**
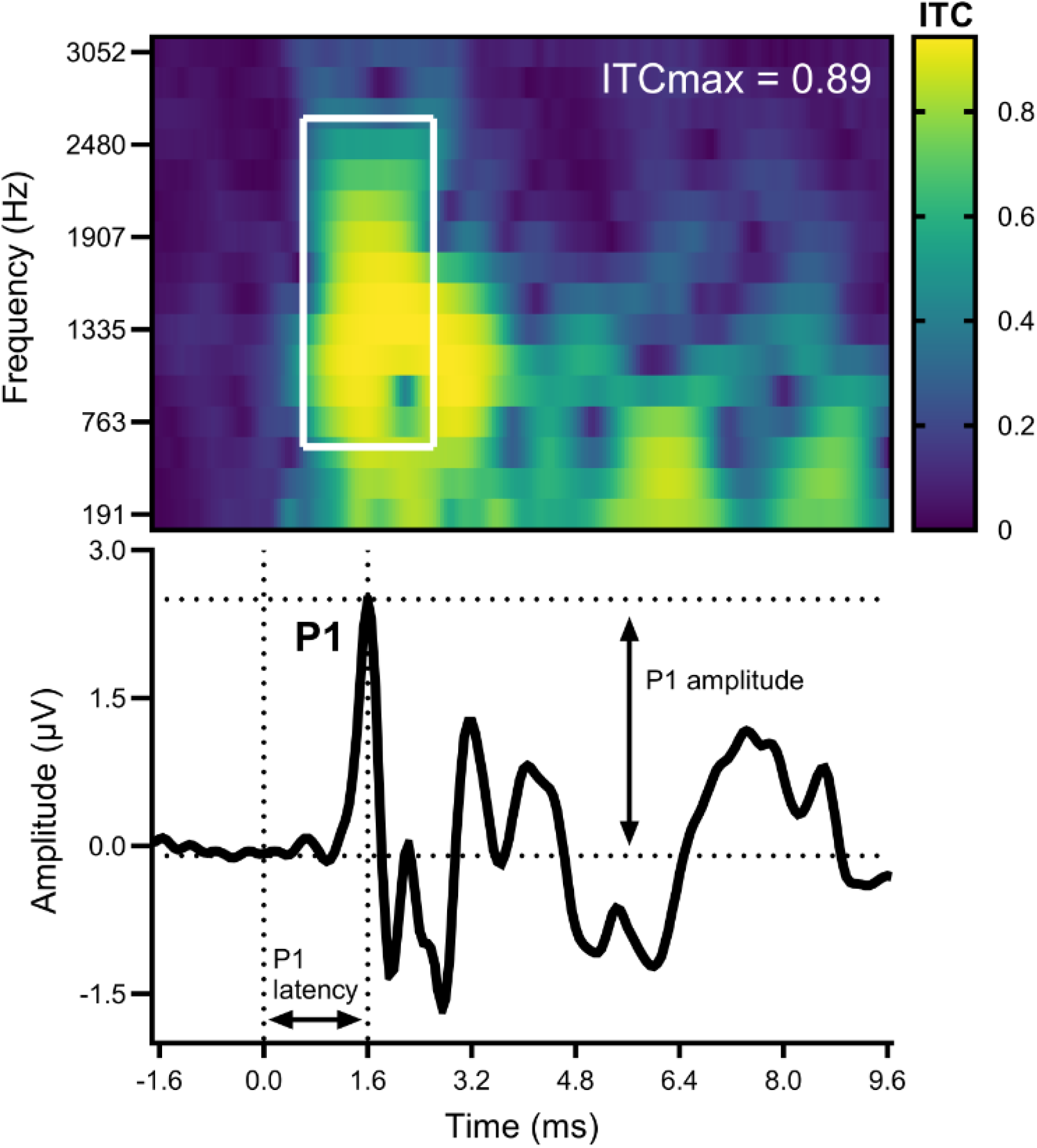
ABR and ITC. The Auditory Brainstem Response (ABR, lower panel) and Inter-Trial Coherence representations obtained from recorded single-trial responses (ITC, upper panel). Both ABR amplitude and ITC in each frequency band are plotted against time (ms). ABR Wave 1 (indicated by P1) amplitude (µV) is measured from the response baseline to the indicated positive peak. ABR Wave 1 latency (ms) is measured from the triggered onset of the stimulus presentation (0 ms) to the time of the indicated positive peak. ITC ranges from 0 to 1, as described in the methods, and is plotted on a color gradient scale from black to yellow, with ITCmax extracted as the maximum value of ITC within a 2 ms time window centred on the wave 1 peak latency and frequency band from 500-2500 Hz, as indicated by the white box.

### Inter-Trial Coherence (ITC)

In addition to looking at the averaged ABR waveform, individual trial-level responses to the 512 stimuli were analysed to measure the coherence of the phase of the response at a set time during these individual stimulus trials. ITC, also called inter-trial phase coherence or phase-locking value, is determined by the coherence between trials of the response phase within each frequency/dB level combination (Lachaux et al 1999). A typical profile of ITC over time and frequency band is plotted in Figure 1 (upper panel), aligned in time with the ABR waveform (Figure 1 lower panel). Regions of high ITC (yellow, arising from high synchrony of neural responses) correlate with the peaks of the ABR, whereas regions of low ITC correlate with regions where there is little or no activity shown in the ABR waveform. The maximum ITC value (ITCmax) in the region around ABR wave 1 was used as an indicator of accurate timing of the hair cell response reflected by synchrony of the auditory nerve response that is measured by wave 1.

Measurements of ABR threshold, ABR wave 1 amplitude, ABR wave 1 latency and ITCmax were standardized to bring the highly differing value ranges of each parameter into a similar minimum-to-maximum range. The standardized values for each parameter are plotted against each other in Figure 2A-F. The standardized values for amplitude and ITCmax (Figure 2E) were subject to k-means clustering, an unsupervised machine learning method. These two metrics were selected because this combination gave the best separation of pathologies, and adding other parameters into the analysis did not add any further discrimination. Suprathreshold response amplitudes are routinely used as a clinical marker. We opted to use a simple two cluster, two parameter k-means analysis to segregate the responses from different mutant mouse lines *a priori* because the main distinction we wanted to find was between two pathologies, a strial dysfunction compared with any other cochlear pathology (Figure 2E; Table 2).

**Table 2.** k-means clustering statistics.

| Cluster | Cluster Size | Silhouette Score | Centroid (X1, X2) | Key Characteristics |
| --- | --- | --- | --- | --- |
| (red) 1 | 444 | 0.6642 | (-0.996, -0.922) | Low X1, Low X2 |
| (blue) 2 | 708 | 0.7123 | (0.625, 0.578) | High X1, High X2 |

**Figure 2.**
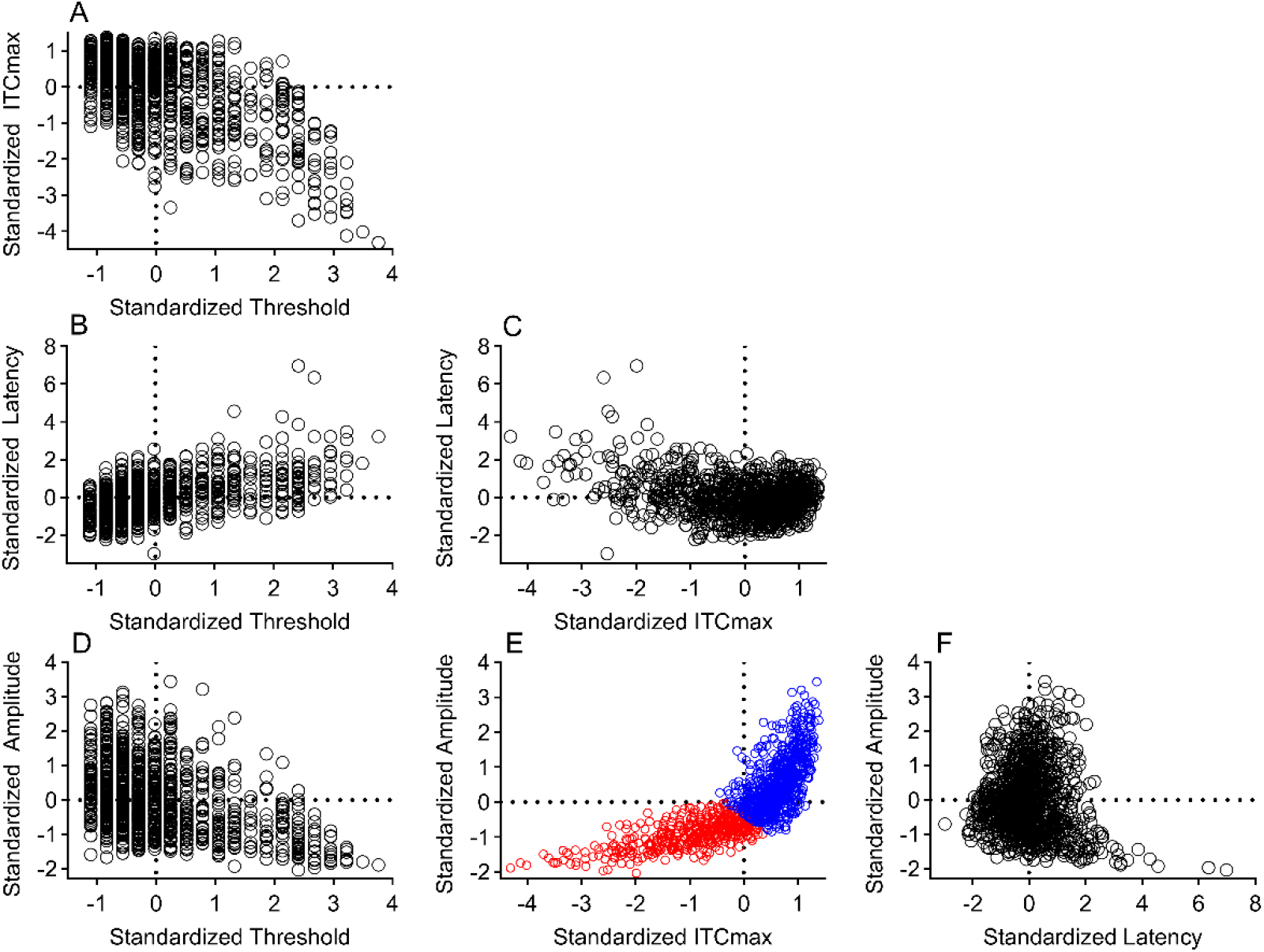
Standardized Parameter Values & k-means clustering. Prior to running k-means clustering, parameter values are standardized. (A) – (F) plot pairs of the parameters measured; ABR threshold, wave 1 amplitude, wave 1 latency and wave 1 ITCmax. Each point represents results from one mouse at one stimulus frequency. (E) Distributions of amplitude and ITCmax specifying 2 clusters in the k-means analysis, with the 2 clusters plotted as red or blue points.

### Distributions of Responses across Pathologies

The arc-like trajectory of values of amplitude and ITCmax are replotted in Figure 3A with points colored according to the mutant. Most mouse/frequency points from littermate control mice show positive amplitude values or positive ITCmax values, lying above the means indicated by the dotted lines (Figure 3B). Young C57BL/6N mice carrying the *Cdh23*^*ahl*^ allele also have mostly positive amplitude or ITCmax values (Figure 3C,D). As the C57BL/6N mice age and develop progressive high-frequency hearing loss, their responses slide downwards and leftwards along the arc trajectory towards lower values (Figure 3E,F). Mutants with primary strial dysfunction, *S1pr2*^*stdf*^ and *Sgms1*^*tm1b*^ homozygotes, have a similar distribution of values as those from control mice (Figure 3G). This is in contrast to the distributions from mutant mice with sensory, neural, or mixed sensory and neural sites-of-lesion, where values were predominantly found in the lower left region of the distribution (Figure 3H-K).

**Figure 3.**
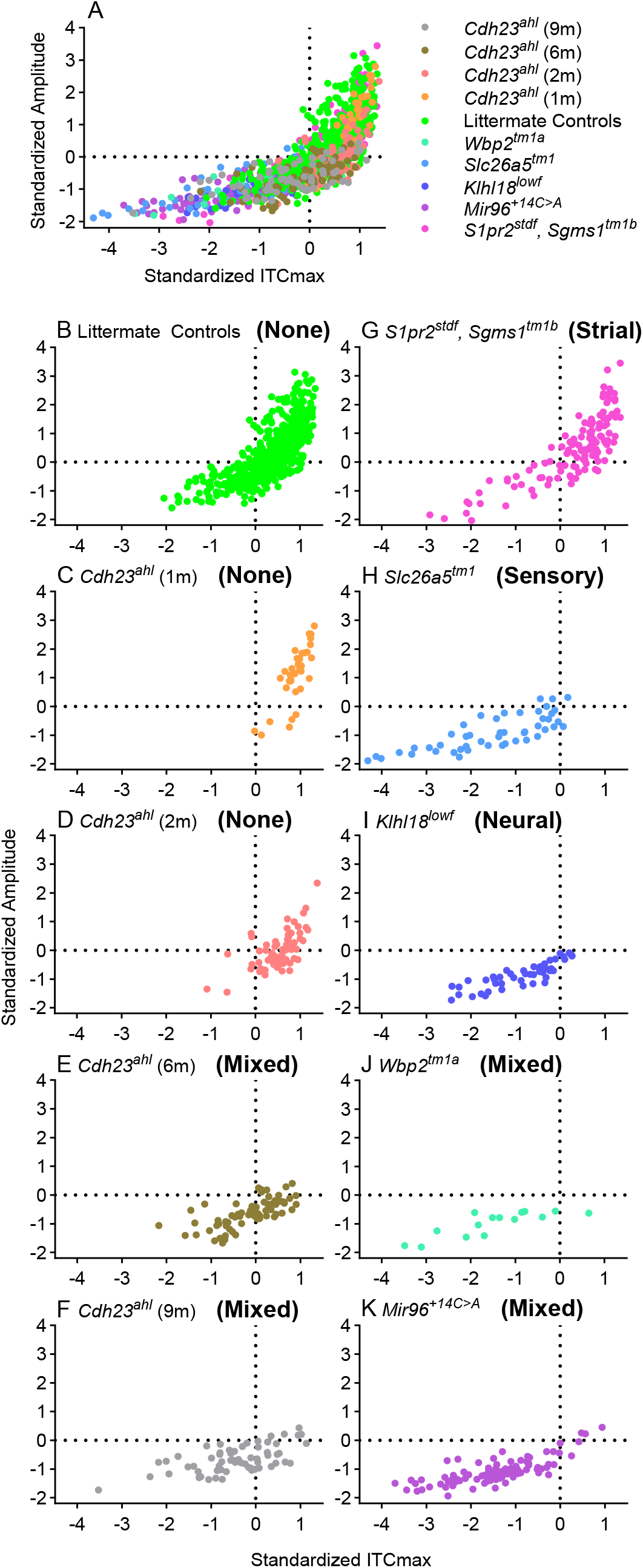
Standardized Parameter Values & Hearing Loss Pathology. (A) The results from Figure 2E are replotted, color coded for the allele (and age) carried by each group. (B)-(F) plot responses from young littermate control mice (B) and from increasing ages of C57BL/6N wildtype mice carrying the *Cdh23*^*ahl*^ allele, from 1 month (C), 2 months (D), 6 months (E) and 9 months (F) old mice. (G)-(K) plot responses from mutant mice with different pathologies. (G) Responses from *S1pr2*^*stdf*^ and *Sgms1*^*tm1b*^ mutant mice with a strial defect. (H) Responses from *Slc26a5*^*tm1*^ mutant mice with an outer hair cell (sensory) deficit. (I) Responses from *Klhl18*^*lowf*^ mutant mice with an inner hair cell (neural) deficit. (J) Responses from *Wbp2*^*tm1a*^ mutant mice with a synaptic defect and reduced DPOAEs (mixed sensory/neural). (K) Responses from *Mir96*^*+14C>A*^ mutant mice with both inner and outer hair cell defects (mixed sensory/neural). Dotted lines represent the means of all values.

### Principal Component Analysis (PCA) and k-means clusters

To provide a more intuitive visual representation of the results, a PCA of the ABR amplitude and ITCmax data was performed (Figure 4A-K), with the 1^st^ principal component plotted on the x-axis (accounting for 88.5% of the variance in the data) and the 2^nd^ principal component plotted on the y-axis (accounting for the 11.5 % of the variance). Boundary lines are also plotted, encompassing responses falling into the two k-means clusters: a blue boundary to represent normal hearing responses and a red boundary to represent impaired responses. For littermate controls and young C57BL/6N mice, 78.0 – 94.4 % of responses fall into the normal hearing cluster (Figure 4B-D). As wildtype mice with the *Cdh23*^*ahl*^ allele age from 1, 2, 6 to 9 months old, the percentage of their responses falling into the normal hearing clusters falls from 94.4%, to 87.0%, to 40.3% and to 28.6%, respectively, as the severity of their mixed sensory & neural pathology progresses (Figure 4C-F). Responses from mice with a strial site-of-lesion also predominantly fall into the normal hearing cluster (77.5%; Figure 4G). Again, this contrast with responses from the other lines of mutant mice with sensory, neural, or mixed sensory and neural pathology (Figure 4H-K), where only 6.3 – 12.4 % fall into the normal hearing cluster. These percentages are also provided in Table 3.

**Table 3.**
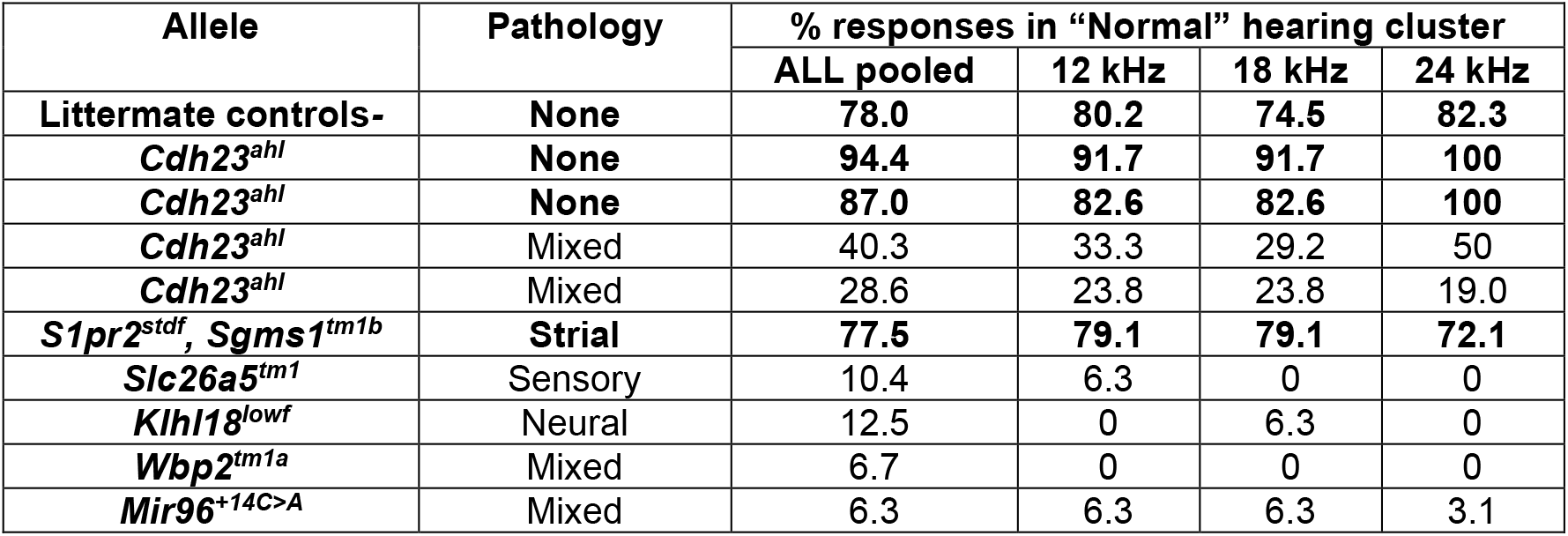
Distribution of ITC responses by pathology in the Normal hearing cluster.

**Figure 4.**
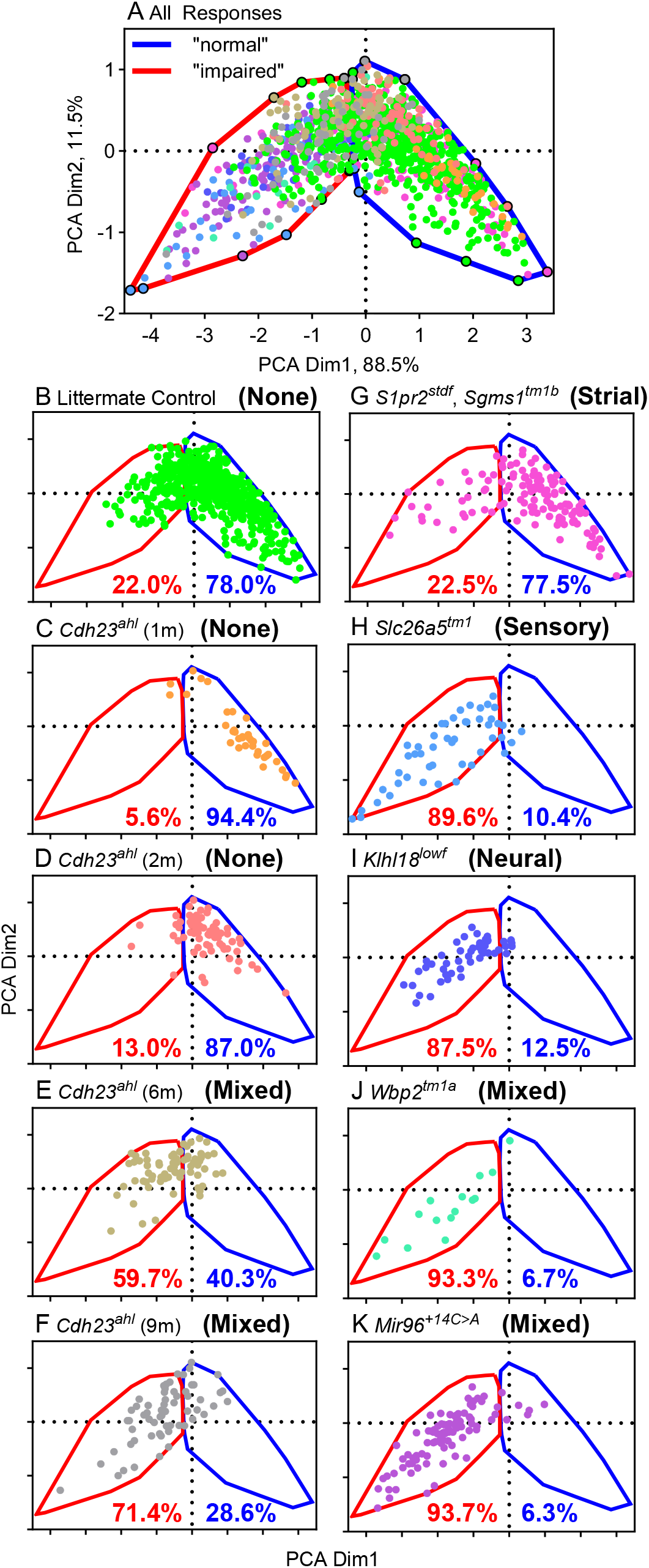
Principle Component Analysis to visualize response segregation between k-means clusters. (A) plots values from PCA dimension 2 against values for PCA dimension 1. Individual points are colour-coded by the different alleles and ages (defined in Figure 3A). The blue and red boundary lines encircle points falling into the blue and red k-means clusters, respectively, as shown in Figure 2E. The blue-bound cluster represents points that come mostly from mice with normal hearing, or from those with a strial deficit. The red-bound cluster represents points that come mostly from mice with a sensory or neural deficit or mixed sensory/neural defect. The remaining panels replot the results from (A), with each experimental group separated onto a separate panel for clarity. (B)-(F) plot responses from young littermate control mice (B) and from increasing ages of C57BL/6N wildtype mice carrying the *Cdh23*^*ahl*^ allele (C-F). (G)-(K) plot responses from mutant mice with different pathologies. The percentage of points falling into the blue or red clusters are indicated on panels (B)-(K) in blue and red text, respectively.

### Relationship between clusters and ABR thresholds

Although our initial analysis of parameters suggested that ABR thresholds did not provide the best separation of sites-of-lesion (see R^2^ values in Figure 2A, B, D), we asked if threshold might contribute further leverage to discriminate between pathology groups. ABR thresholds were plotted separated by the cluster they fell into, normal or impaired hearing (Figure 5). In general, there was considerable overlap in ABR threshold between points from the two clusters, although the points at the highest ABR thresholds tended to be from the hearing-impaired cluster (red points).

**Figure 5.**
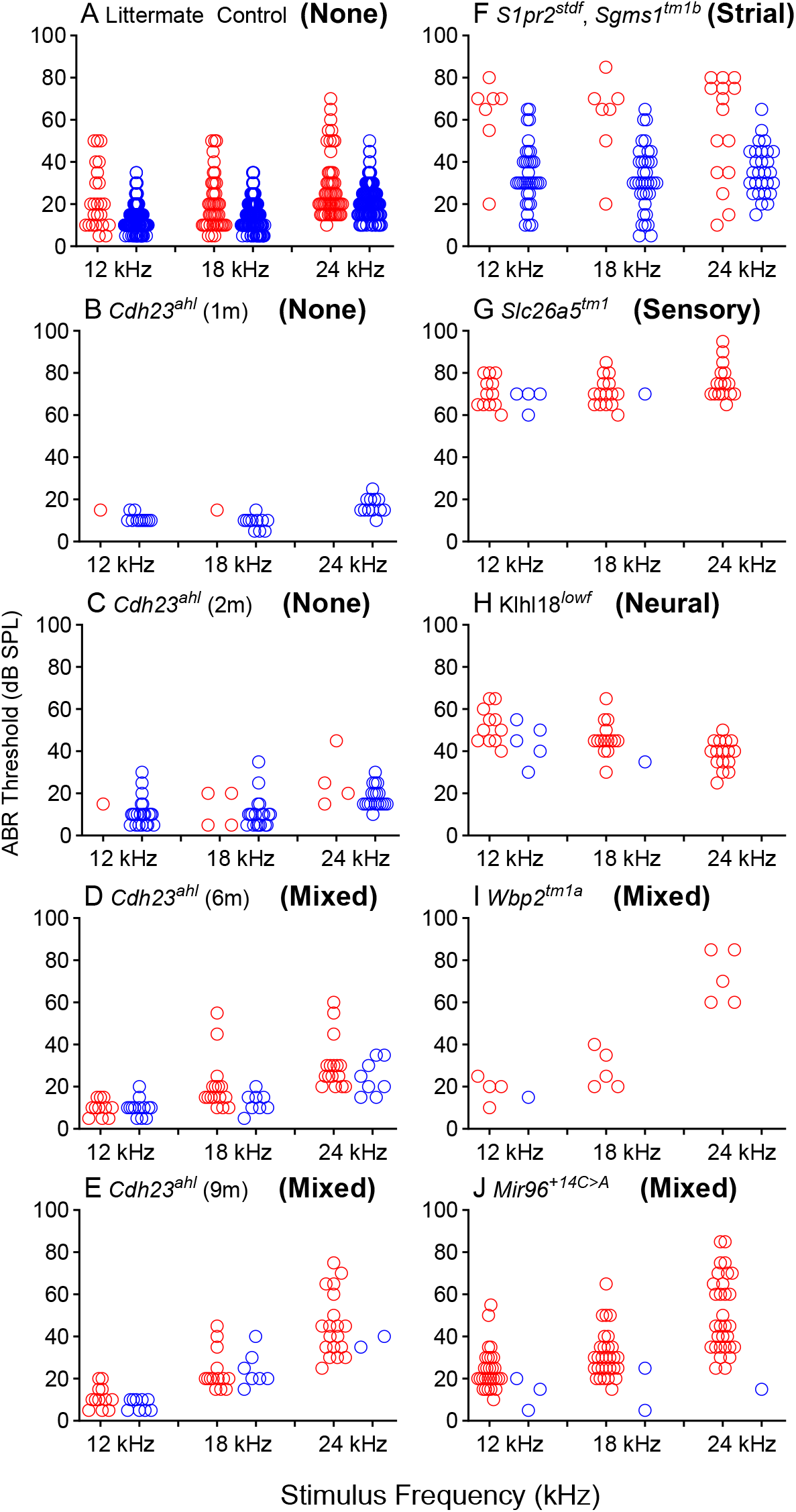
ABR Thresholds. Across the various groups of mice used here (A-J), ABR thresholds are plotted for each stimulus frequency according to the k-means cluster that a particular response was allocated to. Responses in the blue cluster are plotted in blue and those from the red cluster are plotted in red.

## Discussion

We have developed a set of mouse mutants with well-characterised hearing loss due to primary defects at different sites-of-lesion to use in exploring potential non-invasive tests to distinguish reduced strial function from other cochlear pathologies. Using this panel of mice, we have discovered that the ITC/W1amp value can distinguish primary strial dysfunction from primary hair cell or neural defects. In mice with reduced endocochlear potential, the neural responses have a high level of coherence (also known as phase locking value; Harris et al 2018; McClaskey et al 2020) suggesting that hair cells retain their ability to respond to the onset of sound stimuli with high synchrony and accurate timing despite a reduced electrical driving force across their transduction channels and a reduced amplitude of response. Mice with primary hair cell or neural deficits showed reduced ITC/W1amp values. Unsupervised machine learning and clustering of ITCmax values and ABR wave 1 amplitudes placed around 90% of responses from mice with primary sensory or neural defects in the abnormal ITC/W1amp cluster, while only 22% of responses from mice with primary strial dysfunction fell into the abnormal ITC/W1amp cluster. No other measure we used here gave this degree of resolution.

The only other approach currently available to predict strial dysfunction in humans uses the overall shape of the audiogram, which is relatively flat across frequencies in gerbils with reduced EP in contrast to sensory-neural damage that shows steeply sloping thresholds towards high frequencies with low frequencies less affected (Schuknecht 1964; Vaden et al 2022; Kaur et al 2023). We have found a number of mouse mutants that show raised thresholds for ABRs across a wide range of frequencies but with normal EP, including three of the mutants studied in the current panel (*Klhl18*^*lowf*^, *Slc26a5*^*tm1*^, *Mir96*^*+14C>A*^), suggesting alternative approaches will be useful. We appreciate that our panel of mutants may not be representative of the human population.

If strial function is compromised, attempts to treat hair cell or neural deficits by gene therapy or drug treatments are unlikely to be successful, while treatments aimed at restoring strial function may work. We recently found that hearing loss due to a reduced EP could be reversed using a genetic approach to activate the mutant gene (*Spns2*^*tm1a*^) as a proof-of-concept (Martelletti et al 2023), giving optimism that treatments for human hearing loss due to reductions in EP may be successful. However, we cannot measure strial function directly in humans as we can in animals, so a non-invasive diagnostic tool is needed to distinguish between sites-of-lesion. The ITC/W1amp measure was developed in humans before adapting for mouse testing (Harris et al 2018; McClaskey et al 2020; Rumschlag et al 2022; Fabrizio-Stover et al 2025), so it is eminently suitable to use to distinguish strial dysfunction in the clinic. This measure could be used to stratify clinical trials of treatments to maximise the chances of detecting an effect, to inform treatment selection for individuals when these treatments are available, and to guide further development of personalised hearing aids.

## Materials and Methods

### Ethics statement

All mouse work was carried out in accordance with UK Home Office regulations and the UK Animals (Scientific Procedures) Act of 1986 under UK Home Office licences, and the study was approved by the King’s College London Animal Welfare and Ethical Review Body. Both males and females were studied.

### Mouse mutants

The alleles investigated in this study are listed in Table 1 together with the references that describe their phenotypes and origins.

### Physiological Recordings

Single trial Auditory Brainstem Responses (ABRs) were recorded from each mouse under Ketamine/Xylazine anaesthesia (Ingham et al 2011, Ingham 2019, McClaskey et al 2020). Using Tucker Davis Technologies software (BioSigRZ) and hardware (a RZ6 Multi-I/O processor and a Medusa4Z low noise digitizing amplifier), tone pip stimuli (5ms in duration, with a 1ms rise/fall time) of 12, 18 & 24 kHz were presented to the animal at stimulus levels ranging from 0 – 90 dB SPL, at 21 stimuli / sec, and 512 single trial responses recorded (filtered from 100 to 3000 Hz) for a duration of 14ms, epoched from -3 ms to +11 ms, relative to stimulus onset.

### Auditory Brainstem Response (ABR) and Inter-Trial Coherence (ITC/W1amp) Analyses

Using scripts running in Matlab R2023a, the 512 recorded single trial responses recorded for each stimulus frequency and level were averaged to form an ABR (the grand average of the responses). From these ABRs, response threshold was determined and the amplitude and latency of the first positive wave was measured (see Figure 1). Artefact-free single trial responses were analysed via EEGLAB (Delorme & Makeig, 2004; Lopez-Calderon & Luck 2014) to produce an ITC profile for each response. ITC was estimated as the length of the vector formed from averaging complex phase angles from time-frequency decomposition. ITC represents the uniformity of response phase at specific time windows and in specific frequency bands across multiple trials, and ranges from 0 (no phase uniformity) to 1 (perfect phase uniformity across all trials). The maximum ITC value (ITCmax) was obtained from a window 2ms around ABR wave 1 peak (auditory nerve response, indicated by the white box in Figure 1), in a response frequency band from 500-2500 Hz.

### Unsupervised Machine Learning and Cluster Analyses

Values were standardized before further analysis (Xstand = X–mean(X) / sd(X), where X is the parameter value, Xstand is the standardized value, mean(X) & sd(X) are the mean and standard deviation of the values for each parameter). These standardized values were analyzed using k-means clustering (kmeans, Matlab R2023a, using a Squared Euclidean distance metric, 25 replicate initializations with randomised seeds) and Principal Component Analysis (*pca*, Matlab, R2023a). The number of responses falling within each k-means cluster was determined (*inpolygon*, Matlab R2023a).

## Acknowledgments

This work was supported by an International Project Grant from RNID (G86) to NJI & KPS, by Wellcome (221769/Z/20/Z) to KPS and by the National Institute on Deafness and Other Communication Disorders (NIDCD) to KCH. We thank Morag Lewis, Elysia James and Jack Blackburn for mouse colony management and the provision of genotyped mice for these studies.

